# Interrogating Genomic Data in the Phylogenetic Placement of Treeshrews Reveals Potential Sources of Conflict

**DOI:** 10.1101/2021.11.18.469131

**Authors:** Alexander Knyshov, Yana Hrytsenko, Robert Literman, Rachel S. Schwartz

## Abstract

The position of some taxa on the Tree of Life remains controversial despite the increase in genomic data used to infer phylogenies. While analyzing large datasets alleviates stochastic errors, it does not prevent systematic errors in inference, caused by both biological (e.g., incomplete lineage sorting, hybridization) and methodological (e.g., incorrect modeling, erroneous orthology assessments) factors. In this study, we systematically investigated factors that could result in these controversies, using the treeshrew (Scandentia, Mammalia) as a study case. Recent studies have narrowed the phylogenetic position of treeshrews to three competing hypotheses: sister to primates and flying lemurs (Primatomorpha), sister to rodents and lagomorphs (Glires), or sister to a clade comprising all of these. We sampled 50 mammal species including three treeshrews, a selection of taxa from the potential sister groups, and outgroups. Using a large diverse set of loci, we assessed support for the alternative phylogenetic position of treeshrews. The results suggest that the data has statistical support for two hypotheses for the placements of treeshrews, sister to Primatomorpha and to Primatomorpha+Glires. While we observe differences in properties of loci of different types (e.g., CDS, intron, etc.) with respect to the strength of the signal, the support for any particular topology is not dependent on the properties of the data. Rather, we show that the method of phylogenetic signal assessment, as well as whether the signal is measured using the full dataset or only loci with the strongest signal, impact the results much more.

---

In the past two decades, DNA sequence data has become the main source of information for reconstructing phylogenetic relationships. When only a few short gene regions were available for tree inference, stochastic error was often large, resulting in some areas of phylogenies with poor resolution and low branch support. However, the size of phylogenetic datasets has increased dramatically with the advent of High Throughput Sequencing, making it possible to overcome stochastic errors in small datasets and establish some evolutionary relationships with greater certainty (e.g., (Dunn et al. 2008). At the same time, large datasets on their own may still not be able to resolve all phylogenetic relationships due to systematic error (Rodríguez-Ezpeleta et al. 2007). Reanalyses of the same genome-scale datasets with different tools or different data subsets, have resulted in strong support for alternate hypotheses (Chakrabarty et al. 2017; Esselstyn et al. 2017; Reddy et al. 2017). Some relationships remain controversial despite extensive work (e.g. early-diverging animals (Jékely and Budd 2021)], birds (Reddy et al. 2017; Braun et al. 2019)], and placental mammals [Smith et al. 2015, (Murphy et al. 2021)]). Moreover, phylogenetic conflict may be obscured by the inability of traditional support metrics to capture variation in very large datasets, as both non-parametric bootstrap and posterior probability metrics tend to provide strong support for conflicting topologies (Kumar et al. 2012; Salichos and Rokas 2013).

Apparent systematic discordance may be caused by an array of factors. Biological processes such as incomplete lineage sorting (ILS) or hybridization are predicted to be most impactful for areas of the tree of life with rapid diversification (i.e., short internodes) (Rannala et al. 2020). Systematic issues in data processing (such as extensive contamination or homology inference errors) can bias the phylogenetic results of any dataset and are predicted to have a stronger impact on less informative datasets applied to controversial relationships (Simion et al. 2020). Addressing these issues can explain conflicts and resolve phylogenies, as seen in taxa ranging from tunicates to turtles (Brown and Thomson 2017; Simion et al. 2020). Finally, a lack of information in the data may cause essentially arbitrary topological resolution depending on minor factors influencing particular analyses (e.g., (Simion et al. 2020).

In order to disentangle these factors, the array of phylogenetic support methods have evolved to elucidate the presence, strength, and possible cause of phylogenetic conflicts (Simon 2020). For example, the SH-aLRT test (Guindon et al. 2010) assesses likelihood differences between the maximum likelihood (ML) tree and the nearest neighbor interchange (NNI) alternatives for each node, making it a good generic test for informativeness of data. To interrogate locus-specific phylogenetic support, likelihoods (Lee and Hugall 2003; Shen et al. 2017) or marginal likelihoods (Brown and Thomson 2017) can be compared across genes or partitions of interest. This approach provides a gauge of the variation in support among loci and has proven effective in determining the homogeneity of the conflict, and is especially useful at identifying overly impactful outliers (Brown and Thomson 2017; Springer and Gatesy 2018). The site concordance factor (sCF) (Minh et al. 2020a) provides a quartet-based support measure in a parsimony framework, and can be used both on the entire dataset and on a locus-by-locus basis. However, despite the availability of the aforementioned techniques, for many empirical datasets and contentious phylogenetic relationships the extent and potential causes of the conflicts remain elusive.

The position of the treeshrews (Scandentia) is an example of one of the glaring uncertainties in the mammal tree of life (Hallström and Janke 2010; Murphy et al. 2021). Treeshrews are part of the Euarchontoglires, a lineage that also includes rodents, lagomorphs, primates, and colugos. Historically, morphological analyses supported grouping treeshrews with colugos (Dermoptera) (O’Leary et al. 2013); however, genomic data now overwhelmingly support the sister-group relationship between Dermoptera and Primates (Primatomorpha) (Meredith et al. 2011; Esselstyn et al. 2017). One of the currently prevailing hypotheses is that of treeshrews as the sister to the Primatomorpha (Fig. 1) (Hallström and Janke 2008; McCormack et al. 2012; Zoonomia Consortium 2020). This placement, forming the clade Euarchonta, has been recovered by numerous analyses, although typically with relatively low confidence.

**Fig. 1.**
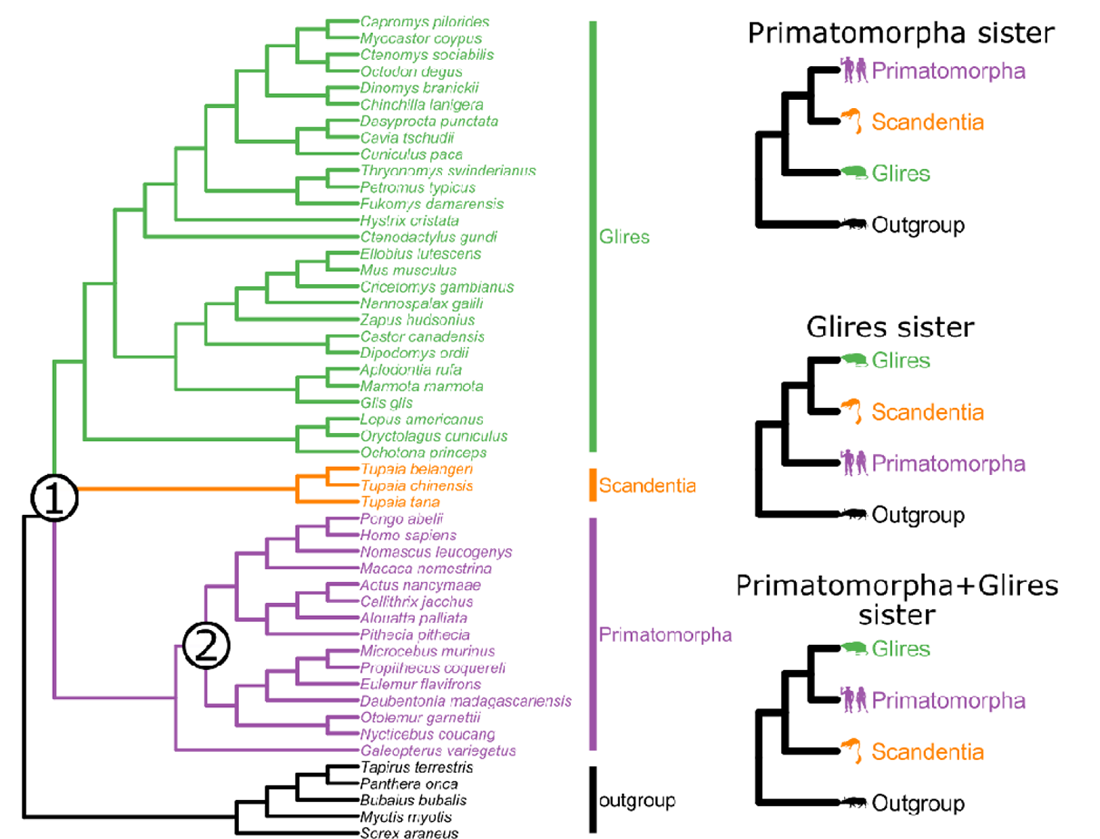
The taxon set and the phylogeny used for evaluations. Different resolutions of node 1 are shown on the right. Node 2 indicates the most recent common ancestor of Primates, this node was used as a control node and was compared to the two alternative nearest neighbor interchanges around the subtending branch.

Alternatively, several phylogenomic analyses have supported the sister-group relationship of treeshrews with rodents and lagomorphs (Glires; Fig. 1) (Tarver et al. 2016; Liu et al. 2017; Du et al. 2019). Finally, one of the most recent molecular analyses (Fig. 1) found that Primatomorpha and Glires themselves form a monophyletic clade, with treeshrews equidistantly related to both (Esselstyn et al. 2017). Analyses recovering each of the topologies have varied in their sampling of species and higher order taxa, the number of sites analyzed, and methodology (Song et al. 2012; Esselstyn et al. 2017; Vachaspati and Warnow 2018), meaning that both biological and methodological factors may be at play when considering the conflicting taxonomic groupings. Thus, with copious genomic data across mammals and a limited set of hypotheses, the placement of treeshrews presents an ideal case study for phylogenetic conflict interrogation (Zoonomia Consortium 2020).

Here we examine the support for the alternative hypotheses of the placement of treeshrews on the mammal tree of life as a case study to understand why we continue to struggle to understand some evolutionary relationships despite genome-scale data. Using millions of variable orthologous sites spread across mammal genomes, we assessed the support for each hypothesis in both parsimony and maximum likelihood frameworks. We observe that the majority of loci are only weakly informative with respect to the phylogenetic placement of treeshrews. The hypothesis receiving the most support depends on both the method of phylogenetic signal assessment and which loci are selected. These results first suggest that treeshrews, Glires, and Primatomorpha might have diversified in rapid succession, and it is challenging to unambiguously distinguish the order of splitting. However, more importantly, the impact of locus selection on phylogenetic results even with very large datasets presents some concern. We suggest that the approach we take here to examine support for alternative hypotheses and gauge the amount of phylogenetic and non-phylogenetic signal, and its potential sources, should be extended to examine other contentious relationships.

## Material and Methods

### Taxon Selection

In order to comprehensively interrogate the conflict in the phylogenetic placement of the treeshrews, we sampled taxa from this group and all major lineages surrounding its most likely position (Primates, Dermoptera, Rodentia, Lagomorpha), as well as five outgroup taxa from Laurasiatheria (Fig. 1). Sampling as many diverse representatives of each group as the publicly available data allow should help to more accurately detect saturating substitutions by breaking long branches and thus improve the signal-to-noise ratio (Baurain et al. 2007; Philippe et al. 2011). Thus we attempted to sample every family with publicly available short read whole genome sequencing data. A total of 50 taxa were included in the dataset (Fig. 1, Table S1).

### Raw Reads

Raw Illumina reads were downloaded from the European Nucleotide Archive FTP server (accession numbers are listed in Table S1). The mean number of paired-end (PE) reads per sample was 955,859,108 (for details see the info in Table S1). Read quality was assessed with FastQC (Andrews 2010).

### Obtaining Orthologous Loci

Putatively orthologous loci were obtained through the SISRS v2.1 pipeline (Schwartz et al. 2015; Harkins et al. 2016; Literman and Schwartz 2021; Literman et al. 2022). Briefly, reads were trimmed using BBDuk (Bushnell 2021) and subsampled using BBMap v38.81 suite scripts (Bushnell 2021) to 10X (0.2X each) assuming an average taxon genome size of 3.5 Gbps. Subsampled reads were assembled in Ray v2.3.1 (Boisvert et al. 2010) to produce a composite genome as the basis of identifying orthologs. The composite assembly was used as a reference for mapping all taxon reads with Bowtie 2 v2.3.5.1 (Langmead and Salzberg 2012; Langmead et al. 2019).

A new step was implemented in SISRS to produce a consensus sequence for each locus (based on the composite genome assembly) for each taxon using the Bowtie 2 aligned data. Sites were called conservatively; sites with depth lower than 3 for the most frequent allele were called as N. Additionally, heterozygous sites were kept track of and consensus sequences with heterozygosity over 1% were discarded, as in the initial tests we empirically determined this to be indicative of improper read alignment. The procedure to produce alignment-based consensus outlined above was chosen over existing solutions (i.e., bcftools consensus), as the latter increases the chances of reference sequence sites propagating into final taxon sequences when coverage or mapping quality are low. This default behavior is undesirable for the SISRS pipeline when analyzing very divergent species, as the reference composite genome sequence is likely to differ significantly from any particular taxon sequence.

Loci were filtered to those containing at least 50% of the taxa, at least one taxon from each of the five major lineages and the outgroup, and to have no more than two-thirds of the sequence missing for each taxon. After filtering, the loci were aligned using MAFFT v7.475 (Katoh and Standley 2013) with the --auto flag.

We filtered potentially misaligned or paralogous data using the Branch Length Correlation (BLC) metric (Simion et al. 2017), which identifies loci whose gene trees contain tip branch lengths least correlated with the concatenation-based tree. Excessively long branches in gene trees may be indicative of alignment or paralogy problems, potentially leading to systematically biased results (Simion et al. 2017). Because the dataset was too large to concatenate all loci, we estimated the concatenation-based tree by analyzing concatenated alignments of 1000 arbitrary loci each in IQ-TREE v2.1.2 with all default parameters except -m GTR+G -bb 1000 -alrt 1000. We then averaged the branch lengths of the obtained trees to produce a set of concatenation-based branch lengths. We reconstructed a gene tree for each locus using IQ-TREE v2.1.2 with all default parameters except -alrt 1000 -m GTR+G (Minh et al. 2020b). We then used the lm function in R to fit the linear model and compute the correlation coefficient, and only retained loci with R-squared more than 0.25.

### Annotation

We hypothesized that locus type (e.g. whether sites fall within coding sequences, introns, etc.) may be associated with the phylogenetic signal for each of the alternative hypotheses, as a similar association has been shown for the overall strength of the signal (Literman and Schwartz 2021). Thus, we annotated the obtained SISRS loci using the human genome as a reference. Human sequences were extracted from the above generated SISRS loci and BLASTN 2.9.0+ (Altschul et al. 1990; Camacho et al. 2009) was used to align these sequences to the human genome assembly GRCh38.p13 (Schneider et al. 2017). A custom script was used to filter the BLASTN output table and convert it into a BED file. Coordinates of the resulting alignments on the human genome chromosomes were intersected (Quinlan and Hall 2010) with the GFF file containing annotations. Another custom script was used to process the annotated BED file and produce a table with a proportion of each type of genomic feature for each locus. We used the following feature types: CDS (coding sequence), UTR (untranslated region, defined as a combination of gene, mRNA, and exon, but no CDS), intron (defined as a combination of gene and mRNA, but no exon), lncRNA (long non-coding RNA), pseudogene, other (other features), and unannotated sequence. For the purposes of subsequent analyses, each SISRS locus was assigned to only one of the types of features based on which type covered the highest proportion of the locus.

### Assessment of Locus Properties

We hypothesized that in addition to the annotation type, other properties of loci may be correlated with the support for particular phylogenetic hypotheses (Mongiardino Koch 2021). QUAST v5.0.2 (Gurevich et al. 2013) was used to assess the basic properties of the assembly, so that their covariation with phylogenetic signal can be investigated. For each locus, we scored the number of taxa with data, locus length, proportion of missing data, the proportion of variable and parsimony informative sites, and GC content using AMAS (Borowiec 2016). The number of taxa and proportion of missing data quantify different levels of missing data, shown to be impactful when non-randomly distributed within phylogenetic datasets (Xi et al. 2016). The locus length and proportions of variable and parsimony informative sites are associated with the number of sites available for model estimates and for resolving relationships (Shen et al. 2016; Steenwyk et al. 2020). Quantifying locus GC content is helpful in determining skewed base composition, which might be indicative of GC-biased gene conversion (Duret and Galtier 2009), which has been shown to impact phylogenetic signal (Romiguier et al. 2013).

Good fit of a substitution model allows more accurate discrimination between synapomorphic and convergent substitutions (Philippe et al. 2011), thus we assessed the absolute fit of a GTR+G model to each locus using PhyloMAd (Duchêne et al. 2017, 2018a, 2018b). The GTR+G model was chosen for all loci as it is one of the most generic substitution models and could be fit to a wide diversity of loci (Sumner et al. 2012; Abadi et al. 2019). As a metric of the adequacy of the model, we used the Mahalanobis distance (Mahalanobis 1936; Drummond and Suchard 2008), which serves as a summary statistic in PhyloMAd. Specifically, we recorded the number of standard deviations of the predictive distribution (SDPD) between the mean of the distribution and the test statistic for the empirical data (Duchêne et al. 2017). The SDPD is expected to be closer to 0 for loci with a good model fit and be further away for loci with a poor model fit. Because minimization of the substitution saturation levels has been shown to improve the phylogenetic signal (Philippe et al. 2011), we assessed saturation in our dataset. Following (Borowiec et al. 2015), we compared corrected (TN93) and uncorrected (raw) pairwise distances in R 4.1.0 (R Core Team 2021) with the package ape 5.5 (Paradis and Schliep 2019). A linear model was fit to the data to estimate the slope of the regression line, with higher values corresponding to lower relative saturation. We performed a principal component analysis on the locus features using R with the ade4 v1.7-17 package (Thioulouse et al. 1997). The results of the analysis were used to assess the correlation between different locus properties and ensure the independence of the chosen metrics.

### Phylogenetic Signal Assessment

To assess overall phylogenetic signal, we first obtained an estimate of the phylogeny based on the complete dataset. We performed a coalescent analysis in SVDQuartets (Chifman and Kubatko 2014) as implemented in PAUP* Version 4.0a, build 168 (Swofford 2003). Only parsimony-informative sites were included and we ran 1000 bootstrap replicates to assess the support for each of the treeshrew placements.

To investigate the phylogenetic signal on a locus-by-locus basis, we used two metrics as proxies for the signal: a log-likelihood difference (Lee and Hugall 2003; Shen et al. 2017) between competing hypotheses (delta log-likelihood, ΔlnL), and a difference in sCF (Minh et al. 2020a) between competing hypotheses. In addition to measuring the phylogenetic signal for the competing hypotheses of the treeshrew placement, we also assessed the signal for the monophyly of primates in order to compare the support for a contentious node with the support for a well-supported split of similar age.

Because assessment of both ΔlnL and sCF required predetermined topologies, we used the mammalian phylogenetic tree of (Armstrong et al. 2020), which was based on the taxa with genomic data available through the Zoonomia project (Zoonomia Consortium 2020) and included 49 of the 50 species we sampled. We pruned the tree to retain only the taxa that we sampled and, since *Tupaia belangeri* was not included in the original tree, we grafted this species as the sister to *Tupaia chinensis* based on (Fan et al. 2013). This topology is well-supported and uncontroversial with regard to the major lineages (Kumar et al. 2017). Because the resulting tree represented only one possible treeshrew placement (Scandentia+Primatomorpha), we obtained the two alternative placements using the phangorn package v2.7.0 (Schliep 2011) in R. These three topologies were used for subsequent evaluations (Fig. 1). To examine the phylogenetic signal for primates, we used the original tree (Fig. 1) with monophyletic primates and two trees with the alternative topologies around the primates branch (i.e. Haplorhini or Strepsirrhini sister to Dermoptera).

Each locus was fit to each phylogenetic hypothesis in IQ-TREE v2.1.2 (Minh et al. 2020b) in two different ways (individually and as part of a larger concatenated alignment) and log-likelihoods were recorded. The approach of fitting loci individually was used to minimize the probability of a single underlying tree assumption violation, which is very likely in the concatenation framework (Rannala et al. 2020). On the other hand, because the loci were on average very short, we hypothesized that the estimates of model parameters may be too inaccurate compared to model estimates for longer alignments (Xia 2020). Thus we also assessed the likelihood-based signal by fitting concatenated partitioned alignments to the alternative topologies, which is a more common approach for conflict interrogation (Shen et al. 2017). Because the total amount of sequence data exceeded the capabilities of the compute cluster system (total length over 15Mbp, see the results), we made the concatenated alignment computationally tractable: loci were arbitrarily binned into groups of 1000, and loci of each group were concatenated. For concatenated alignments we exported per site log-likelihoods and summed them for each included locus according to its coordinates in the alignment (Lee and Hugall 2003; Shen et al. 2017). For both types of likelihood signal assessment we scaled locus log-likelihood by locus length to estimate a mean per site log-likelihood and to avoid bias associated with locus length (Walker et al. 2020).

To assess the overall support across the entire dataset we summed log-likelihoods per locus to produce total log-likelihood for each topology. On a locus by locus basis, the strength of the likelihood support is typically calculated between two competing hypotheses as the difference in log-likelihood (ΔlnL), with positive and negative values representing the support for one or another topology (Shen et al. 2017). Because we considered three competing hypotheses, per site log-likelihood values for each topology were standardized to be at 0 for a hypothesis with the lowest likelihood. Positive values for a given hypothesis thus indicate the ΔlnL-based support for it compared to the least likely hypothesis. We classified loci into three groups, based on topology, based on which loci had the highest log-likelihood. Loci having two best resolutions or equal likelihood between the alternative resolutions were excluded from this assessment.

In addition to assessing the overall distribution of ΔlnL, we separately analyzed the 1000 loci with the largest differences in fit between the alternative topologies, and by extension with a strongest phylogenetic signal (the “outliers” dataset), as well as the full dataset with these loci excluded (the “non-outliers” dataset).

We used IQ-TREE v2.1.2 to calculate sCF (Minh et al. 2020a). For the Scandentia+Primatomorpha and Scandentia+Glires hypotheses we recorded the sCF of the branch subtending the corresponding clades, while for the Scandentia+(Primatomorpha+Glires) hypothesis we recorded the sCF for the branch subtending the Primatomorpha+Glires clade. In addition, we assessed the correlation between sCF and ΔlnL on a locus-by-locus basis to assess similarities between parsimony-based and model-based phylogenetic signals.

We assessed the significance of the difference in total log-likelihood across the dataset for all competing hypotheses using the approximately unbiased (AU) test (Shimodaira 2002) implemented in the R package scaleboot (Shimodaira 2008). The significance of the differences in the numbers of loci favoring each topological resolution was assessed using Pearson’s chi-squared test (Pearson 1900; Agresti 2018), while Kruskal-Wallis One-Way ANOVA (Kruskal and Wallis 1952) followed by pairwise Wilcoxon Rank-Sum Test (Wilcoxon 1945) with Bonferroni correction were used to compare per locus ΔlnL and sCF distributions between the competing topologies.

### Interrelationship Between Phylogenetic Signal and Loci Properties

We tested for a correlation between values of each locus property, and the phylogenetic signal towards particular topological resolutions of the treeshrews. For this question, we analyzed the distribution of ΔlnL in groups of loci favoring each topology, as well as the distribution of previously assessed locus properties in each group. We used Kruskal-Wallis (KW) One-Way ANOVA (Kruskal and Wallis 1952) followed by pairwise Wilcoxon Rank-Sum Test (Wilcoxon 1945) with Bonferroni correction to compare the distributions of the properties of the loci favoring competing hypotheses, except for the locus type proportions, differences in which were assessed using Pearson’s chi-squared test.

All scripts for data analysis are available at: https://github.com/AlexKnyshov/TreeshrewProject

## Results

### Orthologous Loci Generation

On average, 573,949,175 reads per sample (min 90,024,816, max 1,868,772,590, median 576,806,927) passed the trimming and were used for the SISRS pipeline. A total of 11,997,130 SISRS contigs containing 2,208,378,641bp were identified. The largest contig spanned 11,534bp, N50 was 189, L50 was 4,693,728. After filtering based on missing data criteria and the BLC metric, a total of 83,055 loci containing 15,680,010bp were retained for subsequent analyses.

### Properties of Selected Loci

Results of the evaluations are listed in Table S2 and summarized in Table S3 and Figure S1. The number of taxa varied from 25 (minimum passing filter) to 50, with a median of 36. The average locus length was 189bp. Proportions of variable and parsimony informative sites as well as percent missing were nearly normally distributed with means of 0.58, 0.33, and 22% respectively. Mean GC content was 40.3%, although significant differences in GC content were observed between some locus types (Fig. S2). On average, the estimated saturation of the SISRS loci was low, with the mean slope of saturation curve equal to 0.78. The model fit SDPD formed a bimodal distribution, and most loci had a relatively poor model fit (SDPD peak around 9.54), while a smaller group of loci had better model fit (SDPD peak around 5.84). Since the behavior of different test statistics implemented in PhyloMAd is an active area of research (Duchêne et al. 2018a), it is unclear how a poor fit with respect to the Mahalanobis distance-based SPDP relates to the phylogenetic noise. Most features were orthogonal to each other in the PCA, demonstrating a relatively high level of independence, while proportions of variable and parsimony informative characters strongly covaried (Figure S3). Among the loci successfully mapped to the reference and passing the filtering, types of genomic features were annotated in the following proportions: CDS 14.49%, intron 32.84%, lncRNA 11.00%, pseudogene 0.23%, UTR 4.92%, other 0.44%, and unannotated 26.04%.

### Coalescence analysis results

The SVDQuartets inference on all (parsimony-informative) sites yielded a 100% support for the Primatomorpha-sister hypothesis of treeshrew placement and a 100% support for monophyletic primates. Nevertheless, the tree had a Robinson-Foulds distance of 6 (normalized distance of 6.4%) to the reference tree. The differences were restricted to relationships within the outgroup, the placement of Sciuromorpha within rodents, as well as relationships within Lemuriformes in primates. Most of the tree branches had a 100% bootstrap support with the exception of two branches within the outgroup, which received slightly lower support (99.2-99.9%).

### Phylogenetic Signal in the Treeshrew case

*Likelihood-based signal for individually fit loci.—* Summed log-likelihood of all loci individually fit to each treeshrew placement topology was as follows: −117,594,213 for the Primatomorpha sister, −117,594,082 for the Glires sister, and −117,590,171 for the Primatomorpha+Glires sister (Fig. 2a). AU test results indicate that the Primatomorpha+Glires sister hypothesis was significantly more likely (AU test p-value = 0). The number of loci favoring the Primatomorpha+Glires sister hypothesis (N = 19622) was significantly higher (X-squared = 1161.765, df = 2, p-value = 5.32e-253) than the number of loci favoring the other two topologies (N = 14653 and 14047) (Fig. 2c). However, the likelihood-based signal of loci favoring the Primatomorpha sister (median ΔlnL = 0.0039) was significantly higher than that of the Glires sister (median ΔlnL = 0.0034, W = 107379002, p-value = 5.97e-10) or the Primatomorpha+Glires sister (median ΔlnL = 0.0035, W = 151974436, p-value = 3.78e-19) (Fig. 2e). The signal distribution of the top 1000 loci ranked by their ΔlnL values was different, with significantly more loci (X-squared = 98.942, df = 2, p-value = 3.27e-22) supporting the Primatomorpha sister (N = 452) than the Primatomorpha+Glires sister (N = 351) (Figs 2c, 3a). The likelihood-based signal of outlier loci was not significantly different between hypotheses (KW ANOVA, X-squared = 0.12891, p-value = 0.9376) (Figs 2e, 3a). The ΔlnL distributions and counts of favoring loci in the dataset with top 1000 outliers removed were similar to that of the overall dataset, with the Primatomorpha+Glires favoring loci being prevalent (X-squared = 1166.801, df = 2, p-value = 4.29e-254) (Fig. 2c), but the Primatomorpha favoring loci having stronger signal (median ΔlnL = 0.0035, W = 101073540, p-value = 1.68e-04) (Fig. 2e).

**Fig. 2.**
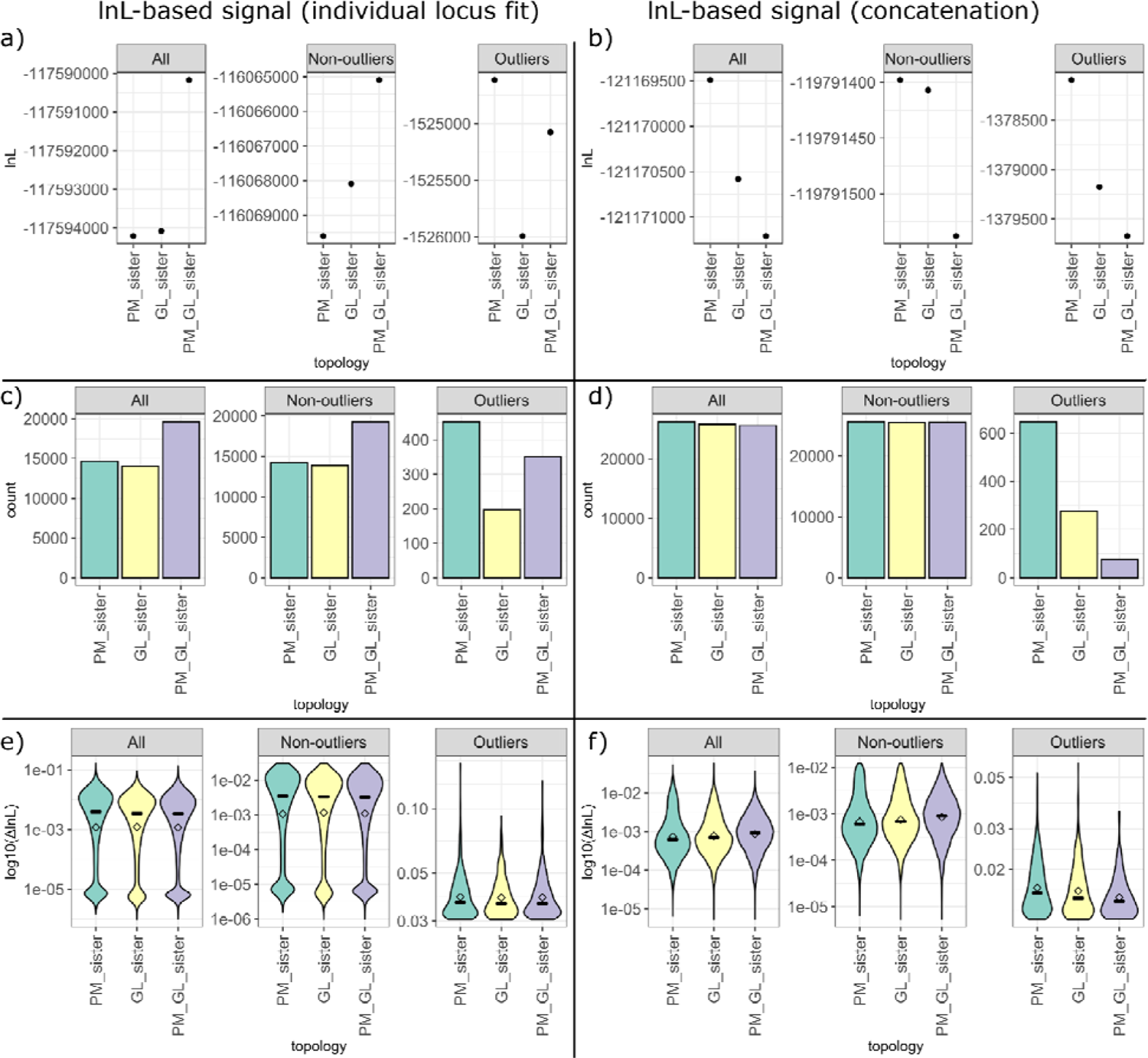
Likelihood-based signal distribution shows ambiguous support for alternative hypotheses of treeshrew placement. The left panel (a, c, and e) shows the results based on loci fit to the trees individually, while the right panel (b, d, and f) shows the same based on loci fit in concatenation, with considerable differences in phylogenetic signal between the two methods of assessment. Each panel features three groups of loci (all, with outliers excluded, and just the outliers), with signal of outliers being different from that of the rest of the dataset. Top row (a and b) shows summed likelihood across all loci for each hypothesis. Middle row (c and d) shows counts of loci that favor (have the highest likelihood) for each of the hypotheses. Bottom row shows the average per-locus likelihood difference between the most and the least likely hypotheses, horizontal lines and points show medians and means respectively.

**Fig. 3.**
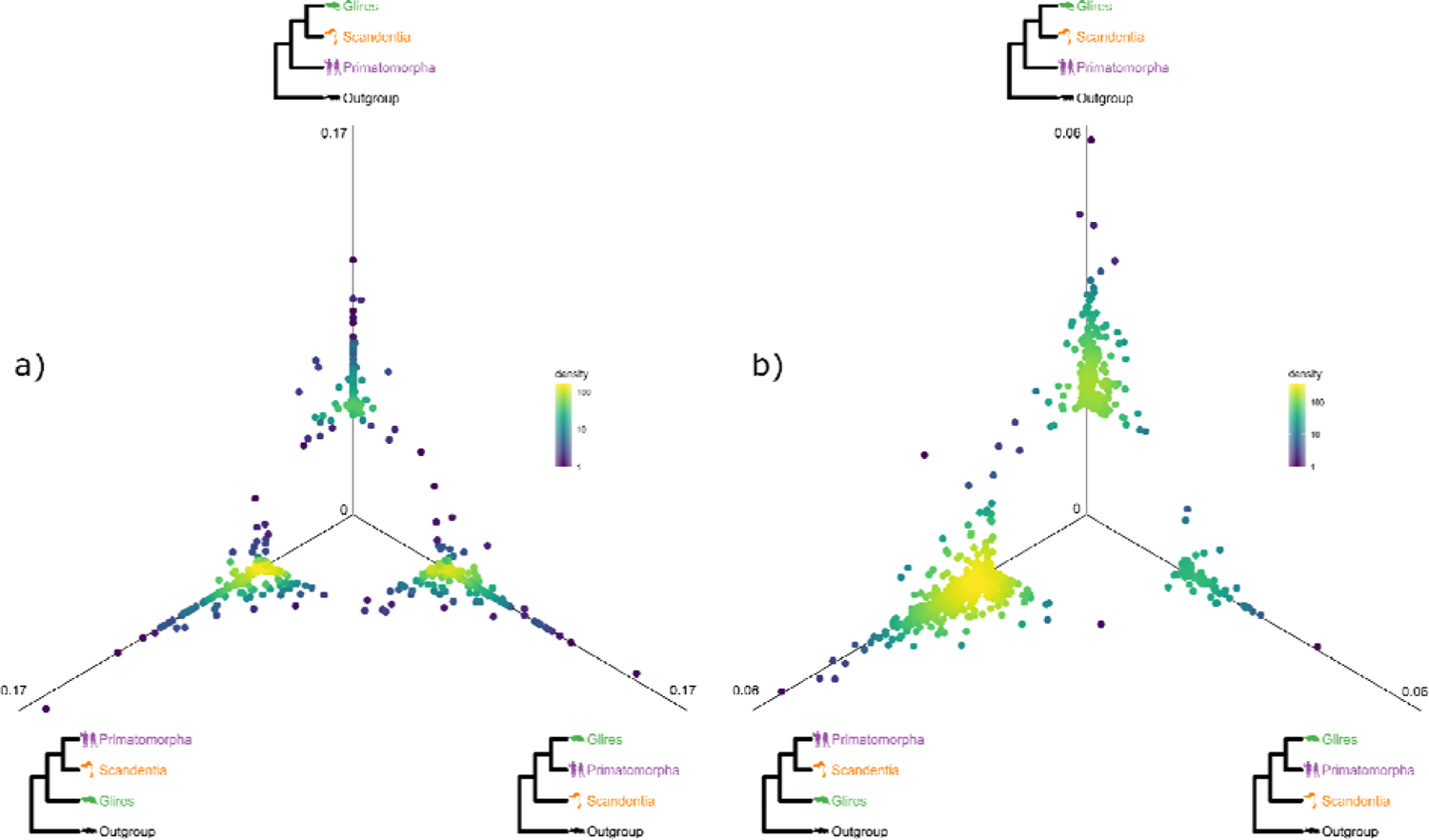
Likelihood support among loci with strong signal (referred to as outliers) for the three competing hypotheses of treeshrew placement, with loci fit individually (a) or in concatenation (b). The figures demonstrate that in both cases more outliers favor the Primatomorpha sister hypothesis, although a noticeable number of loci favor the alternatives. Each panel shows the difference in likelihood (ΔlnL), obtained via fitting each locus to the three competing hypotheses, and rescaled by the locus length. If a point is at the center or close to it, there is no or little difference in likelihood between the topologies for a given locus thus low-likelihood data was omitted. Points plotted further away from the center correspond to loci having higher likelihood under one or two topologies. Each locus point is color coded by density value in that area of the plot.

*Likelihood-based signal for loci fit in concatenation.—* Summed log-likelihood of loci fit to each topology in concatenated alignments was −121,169,491 for Primatomorpha sister, − 121,170,580 for Glires sister, and −121,171,209 for Primatomorpha+Glires sister (Fig. 2b). Thus in concatenation fit the Primatomorpha sister hypothesis was the most likely (AU test p-value = 0.003717603). This was also reflected in the numbers of loci favoring each of the hypotheses: the number of loci favoring the Primatomorpha sister (N = 26261) was significantly higher (X-squared = 7.865172, df = 2, p-value = 0.0196) than the number of loci favoring the alternatives (N = 25808 and 25645) (Fig. 2d). However, the likelihood-based signal strength of loci favoring the Primatomorpha+Glires sister (median ΔlnL = 0.0009) was higher than the signal strength for the alternatives (median ΔlnL = 0.0007 and 0.0006, W = 304381022 and 299618847, and p-values = 1.88e-55 and 2.35e-104 respectively) (Fig. 2f). The signal distribution of the top 1000 loci ranked by their ΔlnL values resembled the overall dataset signal, with significantly more loci (X-squared = 503.34, df = 2, p-value = 5.02e-110) favoring the Primatomorpha sister (N = 647) than the Primatomorpha+Glires sister (N = 76) or the Glires sister (N = 277) (Figs 2d, 3b). The likelihood-based signal strength of outlier loci was not significantly different between the two hypotheses with the highest median signal (W = 97952, p-value = 0.074) (Figs 2f, 3b). The ΔlnL distributions in the dataset with top 1000 outliers removed were similar to that of the overall dataset, with the Primatomorpha+Glires sister loci having stronger signal (median ΔlnL = 0.0009, W = 297286484, p-value = 8.37e-68) (Fig. 2f). However, unlike the results of the dataset with per locus likelihood-based signal, in concatenation there were no significant differences in counts between groups of loci favoring alternative hypotheses when the top 1000 outliers were removed (X-squared = 0.13502, df = 2, p-value = 0.935) (Fig. 2d).

*sCF-based signal.—* Differences in the distribution of per locus sCF in groups of loci favoring alternative treeshrew topologies were significant (KW ANOVA, X-squared = 28.533, p-value = 6.372e-07), however the highest median value was the same for two hypotheses, the Glires sister and the Primatomorpha+Glires sister (median sCF = 40.04 in both groups, W = 347550702, p-value = 4.38e-1) (Fig. 4). Counts of loci favoring each of the alternative treeshrew topologies were also significantly different (X-squared = 328.1939, df = 2, p-value = 5.41e-72), but with more loci (N = 30082) favoring the Primatomorpha sister (Fig. 4).

**Fig. 4.**
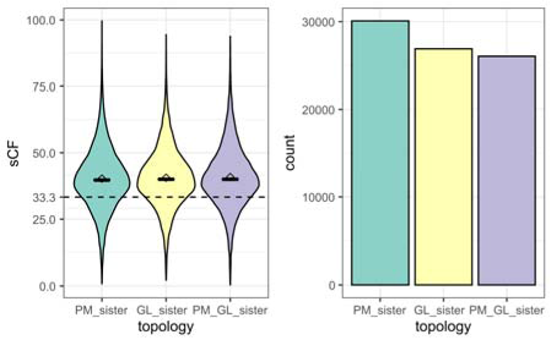
Site concordance factor (sCF) distribution shows weak support for the alternative hypotheses of treeshrew placement, with the Primatomorpha sister having slightly higher counts of loci and all hypotheses having nearly identical median per locus sCF. The left panel shows the distribution of per-locus sCF values, horizontal bars and points show medians and means respectively, and the dashed line shows the majority threshold of sCF (Minh et al 2020). The right panel shows the numbers of loci favoring each topology based on sCF value.

The relationship between ΔlnL and sCF for particular loci was positive but very weak (Fig. S6) both in the per locus fit approach and in the concatenation approach (R-squared 0.07-0.08, p-value < 2.2e-308) (Fig. S4). The same relationship was even more weak for the outlier loci (R-squared 0.008 - 0.03, p-value < 2.2e-308) (Fig. S4).

### Phylogenetic signal in the Primates case

*Likelihood-based signal for individually fit loci.—* Summed log-likelihood of all loci fit individually was −117,594,244 for the control case of monophyletic primates, compared to − 117,695,132 and −117,697,088 for the two alternative NNIs (Fig. 5a). AU test results indicate that primate monophyly was a significantly more likely hypothesis (AU test p-value = 0). The number of loci favoring monophyletic primates (N = 40812) was significantly different (X-squared = 28431.62, df = 2, p-value < 2.2e-308) from the numbers of loci favoring the two alternative topologies (N = 11527 and 10400) (Fig. 5c). The likelihood-based signal of loci favoring monophyletic primates (median ΔlnL = 0.019) was also significantly stronger (W = 316670653, p-value < 2.2e-308) than that of the loci favoring the alternatives (median ΔlnL = 0.006 and 0.007) (Fig. 5e). The signal distribution of the top 1000 loci ranked by their ΔlnL values was different (X-squared = 1934.546, df = 2, p-value < 2.2e-308), with vast majority of loci (N = 989) favoring the monophyletic primates over the two alternatives (N = 6 and 5) (Figs 5c, 6a). The ΔlnL distributions and counts of favoring loci in the dataset with top 1000 outliers removed were similar to that of the overall dataset, favoring the monophyletic primates (ΔlnL: W = 306389180, p-value < 2.2e-308; counts: X-squared = 27021.47, df = 2, p-value < 2.2e-308) (Figs 5d, e). However, the likelihood-based signal of outlier loci was significantly stronger for an alternative topology compared to the monophyletic primates (W = 818, p-value = 0.029), highlighting a possibility for spurious results obtained from likelihood outliers (Figs 5e, 6a).

**Fig. 5.**
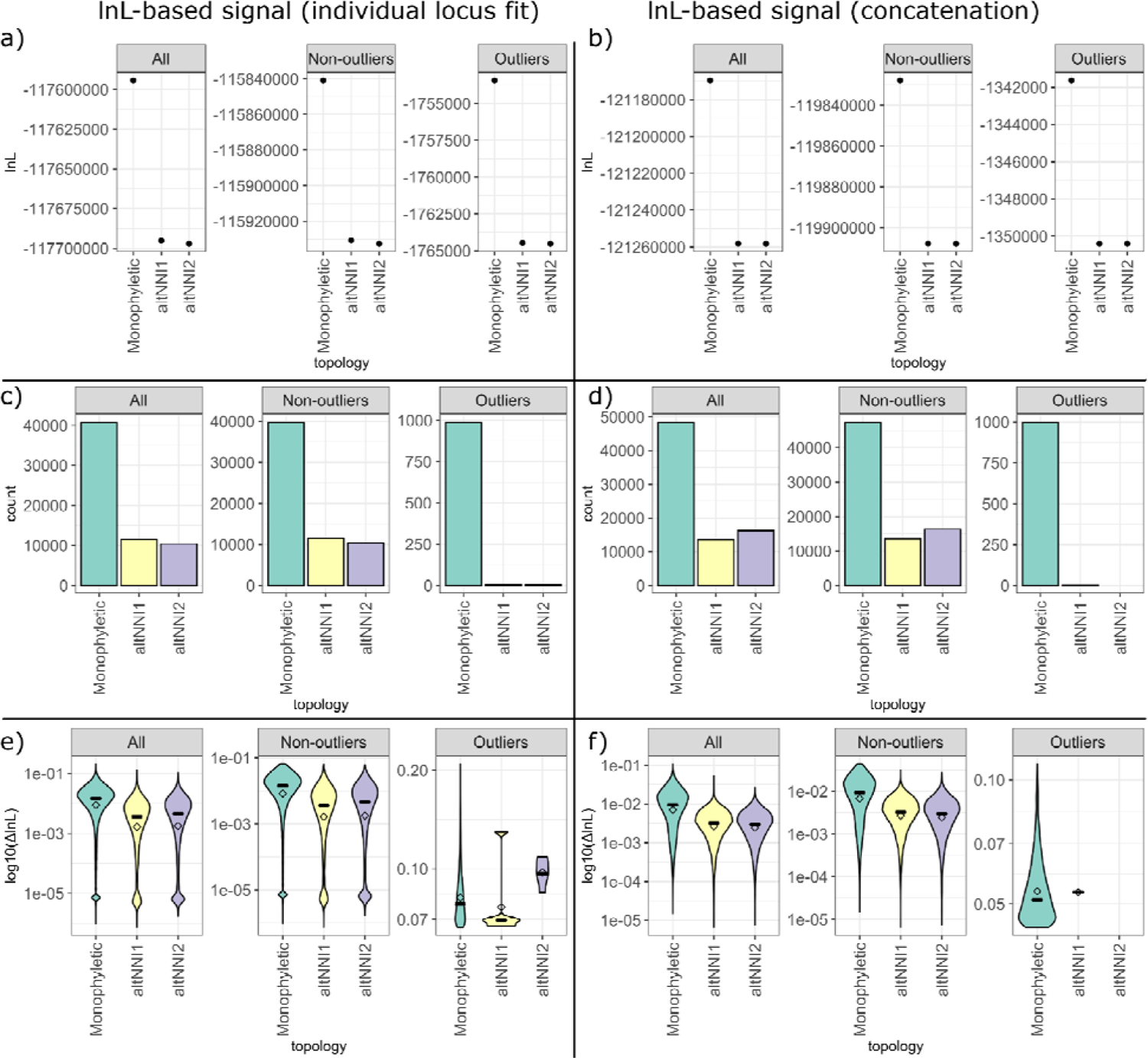
Likelihood-based signal distribution mostly shows strong support for monophyletic primates. The left panel (a, c, and e) shows the results based on loci fit to the trees individually, while the right panel (b, d, and f) shows the same based on loci fit in concatenation, with minimal differences in phylogenetic signal between the two methods of assessment. Each panel features three groups of loci (all, with outliers excluded, and just the outliers), with almost all outliers favoring monophyletic primates. Top row (a and b) shows summed likelihood across all loci for each hypothesis. Middle row (c and d) shows counts of loci that favor (have the highest likelihood) for each of the hypotheses. Bottom row shows the average per-locus likelihood difference between the most and the least likely hypotheses, horizontal lines and points show medians and means respectively.

**Fig. 6.**
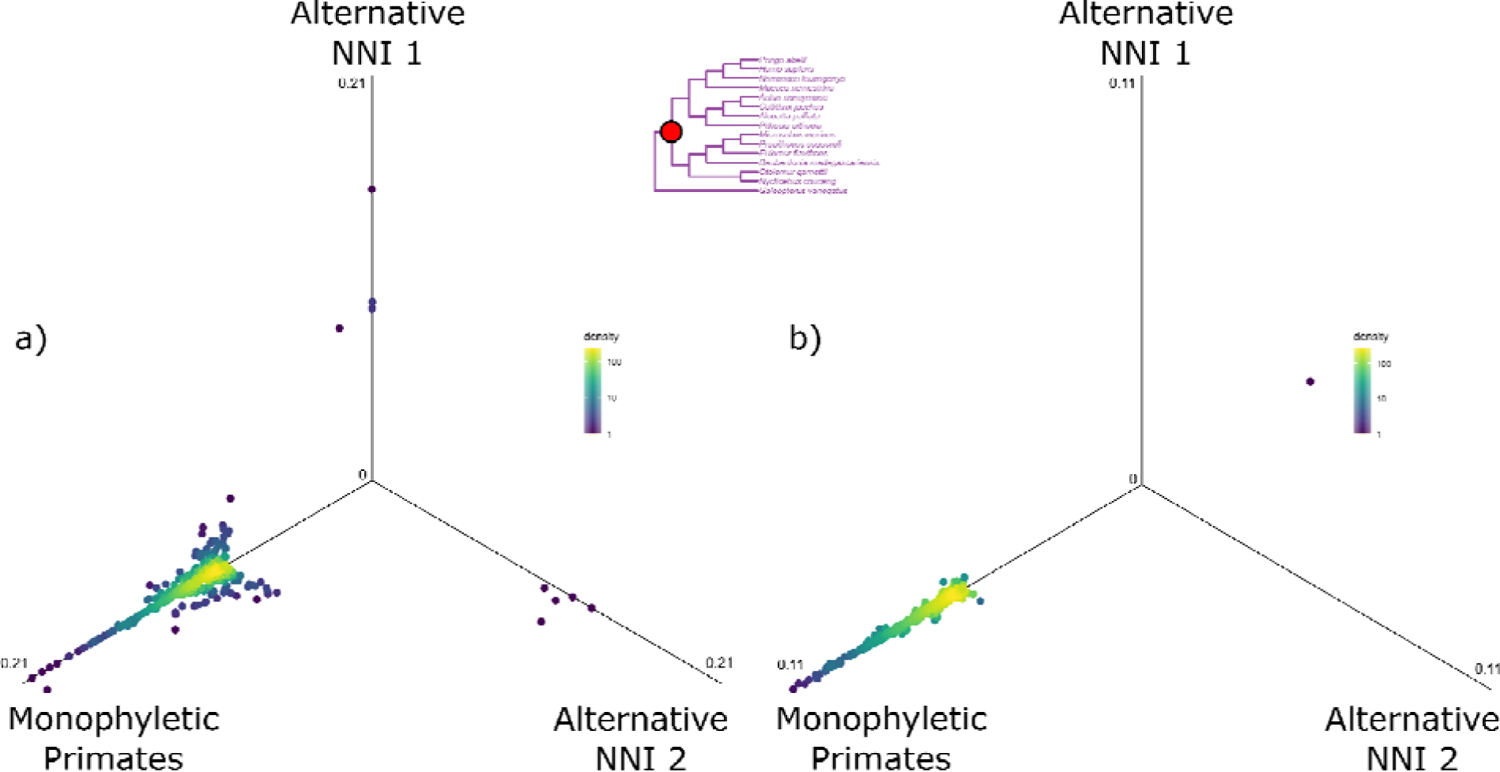
Likelihood support among outliers for monophyletic primates, with loci fit individually (a) or in concatenation (b). In both cases the majority of outliers favor the monophyly as opposed to the alternative hypotheses. Each panel shows the difference in likelihood (ΔlnL), obtained via fitting each locus to the three competing hypotheses, and rescaled by the locus length. If a point is at the center or close to it, there is no or little difference in likelihood between the topologies for a given locus thus low-likelihood data was omitted. Points plotted further away from the center correspond to loci having higher likelihood under one or two topologies. Each locus point is color coded by density value in that area of the plot. The subtree in the center shows the location of th node for which the alternative hypotheses were considered.

*Likelihood-based signal for loci fit in concatenation.—* Summed log-likelihood of all loci in concatenation was −121,169,473 for the monophyletic primates, compared to −121,258,158 and −121,258,201 for the two alternative NNIs (Fig. 5b). AU test results indicate that primates monophyly was a significantly more likely hypothesis (AU p-value = 0). The number of loci favoring monophyletic primates (N = 48412) was significantly higher (X-squared = 28892.28, df = 2, p-value < 2.2e-308) than the numbers of loci favoring the two alternative topologies (N = 13531 and 16242) (Fig. 5d). The strength of the likelihood-based signal of loci favoring monophyletic primates (median ΔlnL = 0.0095) was also significantly higher (W = 495031550, p-value < 2.2e-308) than that of the loci favoring the alternatives (median ΔlnL = 0.0032 and 0.0030) (Fig. 5f). The signal distribution of the top 1000 loci ranked by their ΔlnL values was different (X-squared = 1994.006, df = 2, p-value < 2.2e-308), with all except one locus favoring the monophyletic primates over the two alternatives (Figs 5d, 6b). The likelihood-based signal of outlier loci was not significantly different between hypotheses (KW ANOVA, X-squared = 0.096123, p-value = 0.7565) (Figs 5f, 6b). The ΔlnL distributions and counts of favoring loci in the dataset with top 1000 outliers removed were similar to that of the overall dataset, favoring the monophyletic primates (ΔlnL: W = 481514670, p-value < 2.2e-308; counts: X-squared = 27557.74, df = 2, p-value < 2.2e-308) (Figs 5d, f).

*sCF-based signal.—* Differences in the distribution of per locus sCF between the monophyletic primates and two alternative NNIs were significant (W = 540094683, p-value < 2.2e-308), with the sCF-based signal of the loci favoring monophyletic primates (median sCF = 49.78) being stronger than that of loci favoring two alternative NNIs (median sCF = 41.22 and 40.69) (Fig. 7). The median sCF support for the monophyletic primates was higher than that for any of the treeshrew topologies. Counts of loci favoring the monophyletic primates and alternative topologies were also significantly different (X-squared = 15976.57, df = 2, p-value < 2.2e-308), with the majority of loci (N = 44549) supporting primate monophyly (Fig. 7). Thus, sCF based phylogenetic signal supported primates monophyly much more strongly than any of the competing treeshrew topologies.

**Fig. 7.**
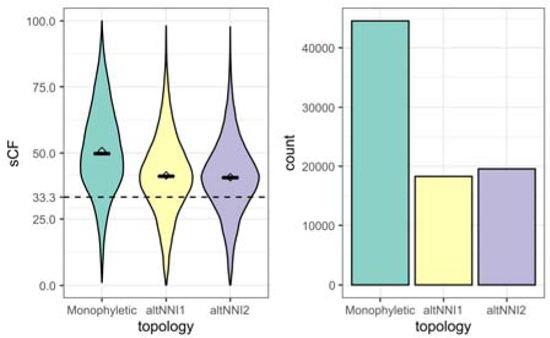
Site concordance factor distribution shows strong support for monophyletic primates. The left panel shows the distribution of per-locus sCF values, horizontal bars and points show medians and means respectively, and the dashed line shows the majority threshold of sCF (Minh et al 2020). The right panel shows the numbers of loci favoring each topology based on sCF value.

The relationship between ΔlnL and sCF across all loci for the primates case (standardized slope 0.43-0.47, R-squared 0.19-0.22, p-value < 2.2e-308) was stronger than in the case of competing treeshrew topologies (Fig. S5). The same relationship for the outliers was weak (R-squared 0.008-0.03, p-value < 2.2e-308) (Fig. S5).

### Support for Treeshrew Relationships Using Loci Supporting Monophyletic Primates

Only 0.6 to 0.9% of top 1000 outlier loci overlapped between the Treeshrew and the Primates analyses, suggesting that the signal in loci varies among nodes. When fit individually, both loci that supported monophyly of primates and those that contradicted statistically favored the Primatomorpha+Glires placement of treeshrew (X-squared = 802.4895 and 141.5346, p-value = 5.52e-175 and 1.85e-31 respectively). Specifically, 42% of loci that favor primates’ monophyly favored the dominant Primatomorpha+Glires hypothesis, while only 38% of loci that did not favor primates monophyly supported that treeshrew placement.

When fit in concatenation, the non-outlier loci that favor monophyletic primates loci favored three competing hypotheses in roughly the same proportions (∼33%, X-squared = 3.904185, df = 2, p-value = 0.142), similarly to the overall signal of non-outliers in concatenation (see above). However, loci that rejected monophyly of primates had a slightly higher but nevertheless significant proportion of support for the Primatomorpha sister (34%, X-squared = 9.281649, df = 2, p-value = 0.00965).

### Interrelationship Between Phylogenetic Signal and Loci Properties

We hypothesized that annotation and other properties of loci may be associated with the signal for each of the competing topologies. When fit individually, loci favoring each of the treeshrew placements had different distributions of most of the properties we assessed, specifically between the loci favoring the Primatomorpha+Glires-sister and the other alternatives (with the exception no differences in alignment length and model fit) (Fig. 8, Table S3). When fit in concatenation, similarly, most properties had significant differences between treeshrew placement topologies, although there was a less clear pattern of which properties were statistically different between which hypotheses (Fig. 9, Table S3).

**Fig. 8.**
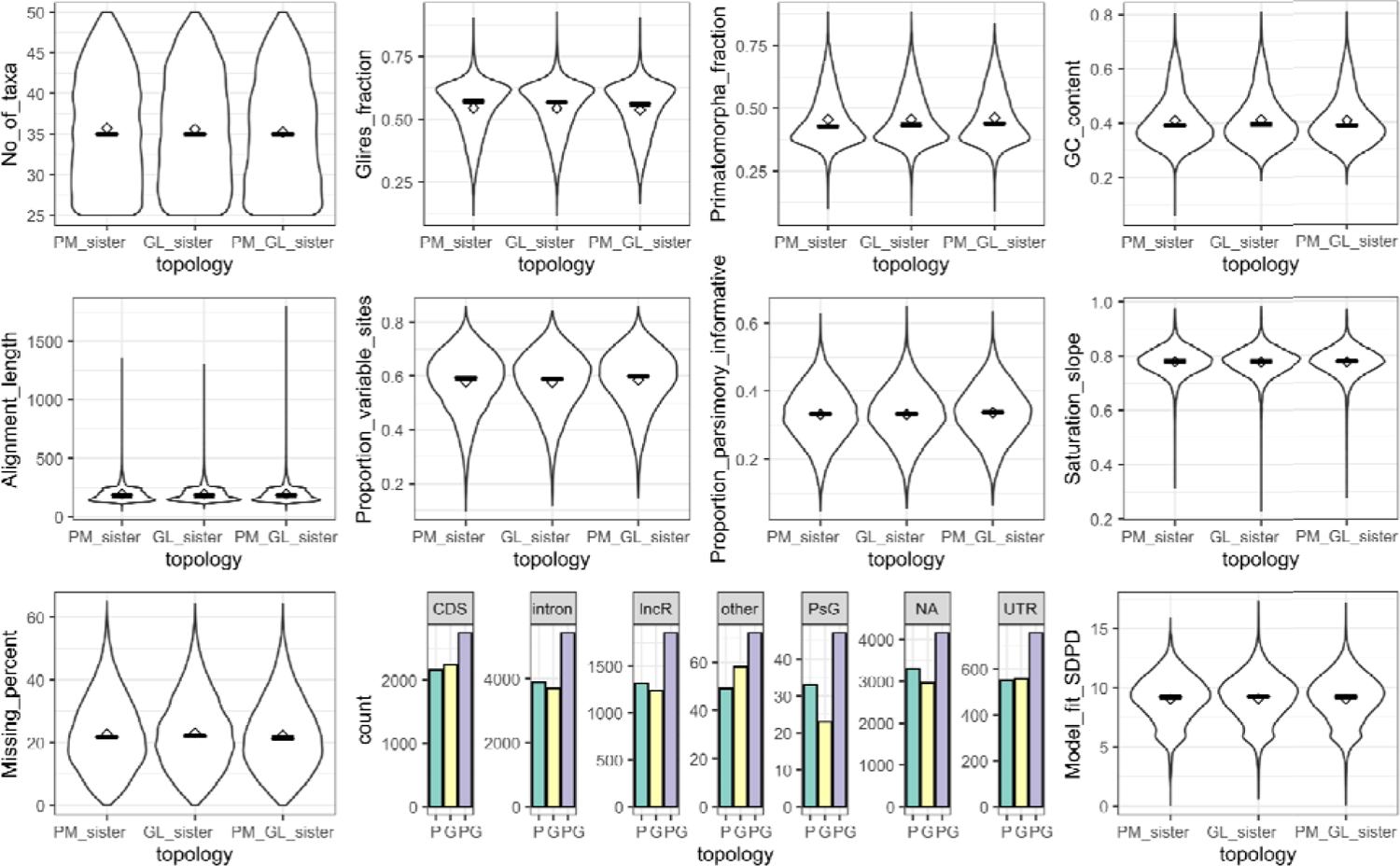
Properties of the loci favoring each of the treeshrew phylogenetic hypotheses when loci fit to alignments individually. Most properties have distributions that look very similar between competing treeshrew topologies. Nevertheless distributions of all properties except for alignment length and model fit SDPD have statistically significant differences. The Primatomorpha+Glires-sister hypothesis is the most frequently supported across all types of loci.

**Fig. 9.**
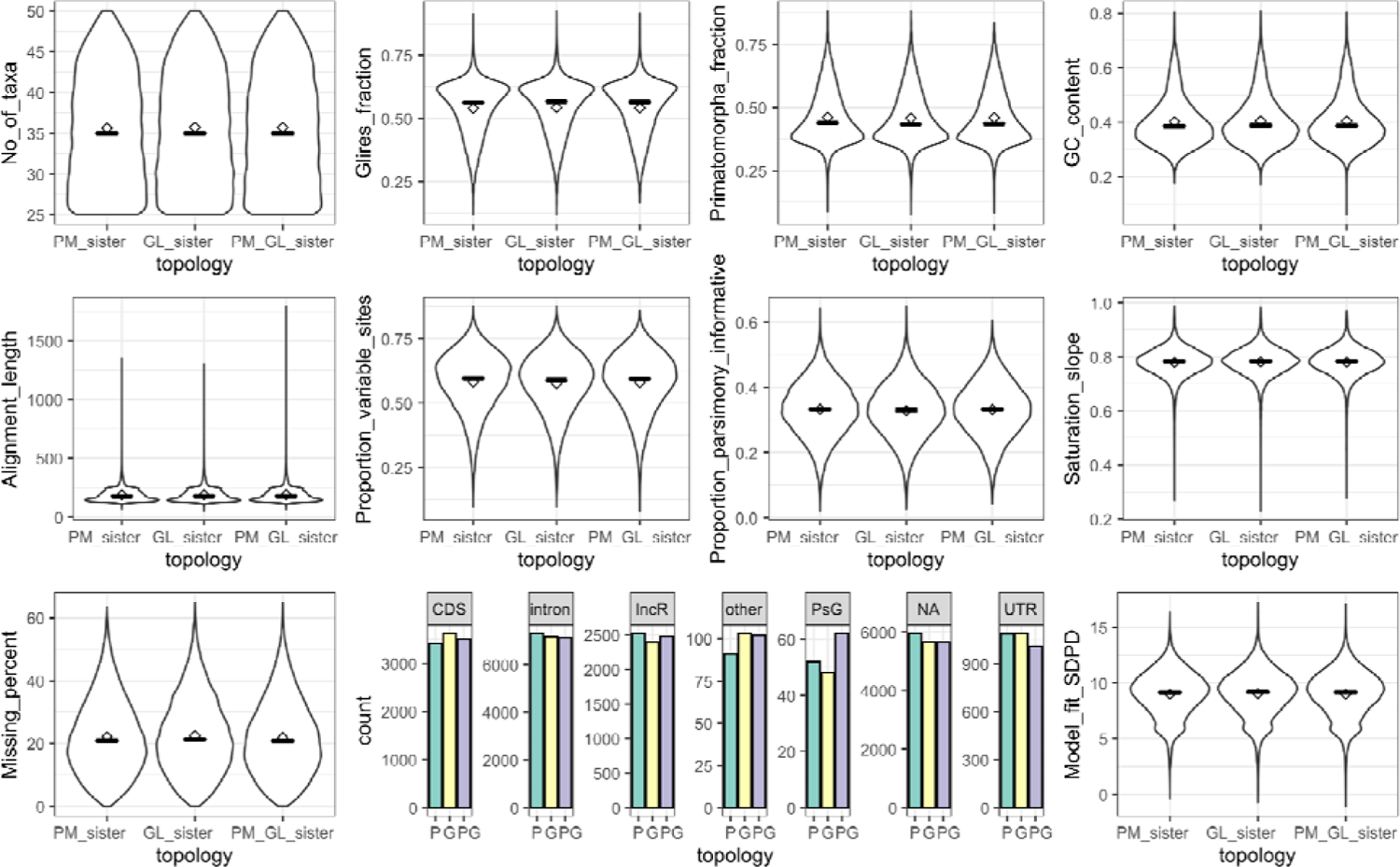
Properties of the loci favoring each of the treeshrew phylogenetic hypotheses when loci fit to alignments in concatenation. Most properties have distributions that look very similar between competing treeshrew topologies. Nevertheless distributions of all properties except for the number of taxa have statistically significant differences between at least two hypotheses. Only among the unannotated loci there were statistically significant prevalence of the Primatomorpha-sister favoring loci.

The proportions of locus types in loci favoring all competing treeshrew topologies were significant when loci were fit individually, while only the proportions between the Primatomorpha-sister and Primatomorpha+Glires-sister were significantly different when loci were fit in concatenation. The counts of loci favoring each treeshrew topology in each locus type overall reflected the total dataset counts. When fit individually, most locus types favored the Primatomorpha+Glires sister (with exception of “other” where the difference was not significant, primarily due to small sample size) (Fig. 8). When fit in concatenation, only the “unannotated” group had significant differences between the competing treeshrew topologies, with other locus types having statistically indistinguishable counts of loci supporting the alternatives (Fig. 9).

## Discussion

Overall, the SISRS loci contained strong phylogenetic signal, as measured on an uncontroversial split (Primates). Both metrics used as proxies for phylogenetic signal - ΔlnL and sCF - showed that the overall dataset clearly favored the topology with monophyletic primates over the two alternative NNIs. With respect to phylogenetic placement of treeshrews, despite observing a strong support for the Primatomorpha-sister hypothesis in the coalescent analysis of all parsimony informative sites, we found disagreements among different proxies for per-locus phylogenetic signal. Additionally, as opposed to the Primates split case, the overall modest per-site ΔlnL-based signal suggests that the majority of genomic markers contain only weak phylogenetic signal for short branch splits, and some contain non-phylogenetic signal.

### Phylogenetic Signal, the Competing Treeshrew Hypotheses, and Primate Monophyly

Overall, two topologies (Primatomorpha-sister and Primatomorpha+Glires-sister) were supported across different analyses. Differences in which particular hypothesis was supported were observed between likelihood-based and site concordance-based analyses, between two types of likelihood-based signal assessment, whether selection was based on average per-locus signal of loci or numbers of loci, as well as whether the entire dataset, only likelihood-based outliers, or the dataset without outliers were considered (Fig. 10). The sCF based phylogenetic signal was also ambiguous with respect to placement of treeshrews.

**Fig. 10.**
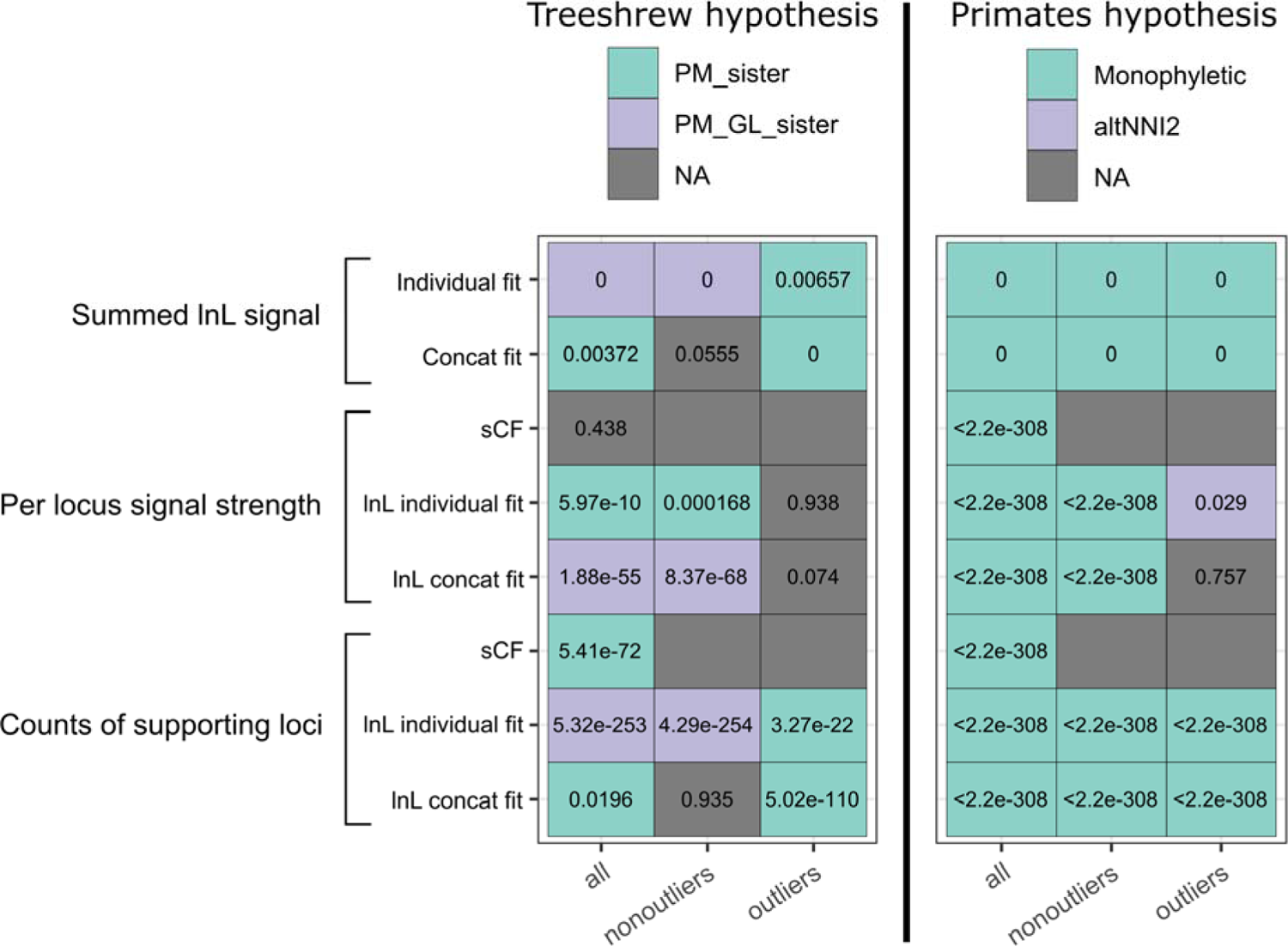
Phylogenetic signal for competing hypotheses summarized over all metrics, highlighting conflicting phylogenetic signal with respect to a contentious node (treeshrew placement) while showing a very strong support for an uncontroversial split (primates monophyly). Color encodes the hypothesis supported by each metric in each subset, gray represents no significant difference between the top two or all three hypotheses, values in the cells are p-values.

When the likelihood-based signal was assessed individually per locus, the total signal (summed lnL across all loci) as well as counts of supporting loci favored the Primatomorpha+Glires-sister. However, per locus signal strength was higher in the group of loci favoring the Primatomorpha-sister. Additionally, the total signal and counts in the strong signal outliers subset also supported the Primatomorpha-sister. When the likelihood-based signal was assessed in concatenation, the signal distribution changed: the total signal and counts were unequivocal except for when the strong signal outliers were included, while per locus signal strength was higher in the group of loci favoring the Primatomorpha+Glires-sister. In both types of likelihood assessment we observed a discordance between loci that have stronger signal and loci that have weaker signal (compare per locus signal strength and locus counts on the figure). With respect to the sCF-based signal, the majority of the loci favored the Primatomorpha-sister, although sCF-based signal strength for that hypothesis was not significantly higher than for the others. Thus, despite that the results of coalescent analysis convey superficial uniformity of the support for the Primatomorpha-sister hypothesis, there is a significant conflict between individual loci in the dataset. Given that, we conclude that based on our data it is impossible to unequivocally resolve treeshrew position.

The overall ambiguity of the signal with respect to the treeshrew placement was considerably larger than for the case of primate monophyly, which is consistent with a notion of rapid radiation between the major Euarchontoglires lineages (Zhou et al. 2015; Esselstyn et al. 2017). However, even for primate monophyly the support was never maximal (all loci), thus showing detectable conflict even in the uncontroversial split. Similar results were found by (Esselstyn et al. 2017) when examining the likelihood support for a well-established monophyly of Glires. We thus argue that non-phylogenetic signal is noticeable when examining both the well supported and the poorly supported nodes, however in the case of the poorly supported one the relatively small amount of phylogenetic signal prevents us from unambiguously resolving the topology.

Locus support (or lack of) for monophyletic primates did not impact the signal distribution with respect to competing treeshrew topologies. Loci that favor monophyly of primates constitute about two thirds of the overall dataset depending on the method of signal assessment, and the distribution of the signal for treeshrew placement in this subset replicated that in the overall (unfiltered) dataset. Thus based on our results it can be concluded that filtering loci based on the support for an uncontroversial relationship would not necessarily improve the phylogenetic signal to help resolve a contentious node.

### Impact of Individual vs in Concatenation Assessment on Phylogenetic Signal

Our results show stark contrast between the two types of assessment of phylogenetic signal, by fitting locus alignments to topologies individually, or as a part of concatenated alignments. We hypothesize that poor model fit for short single-locus alignments may be one reason why the results of likelihood assessments between the two methods differ (Xia 2020). The SISRS loci in our dataset had a mean length of about 200bp, which for conserved loci is likely insufficient to adequately estimate substitution model parameters. Thus the parameter estimates were likely to have random noise, which might increase the likelihood for the alternative topologies and decrease the likelihood for the true tree. The use of short loci has been criticized previously on the grounds of poor model fit and resulting poor gene tree estimates (Bryant and Hahn 2020; Rannala et al. 2020). While normally model choice and fit does not impact the resulting topology as much as it impacts branch length estimates, model accuracy could be critical for nodes that are difficult to resolve (Kelchner and Thomas 2007). This is well illustrated in the case of the order of divergence of Porifera and Ctenophora, where using different types of models results in reconstruction of two alternative topological hypotheses (Li et al. 2021).

However, it is also possible that model parameters may be estimated incorrectly due to concatenating loci. Model parameter estimation for concatenated datasets is commonly done with the help of the tree reconstructed on the entire matrix (in the case of IQ-TREE, it is a parsimony-based tree (Minh et al. 2020b)]). Such a procedure would violate the assumption of multispecies coalescence (MSC) and potentially bias individual loci to support the predominant topology, which they would otherwise reject (Walker et al. 2020). Longer concatenated loci have been noted as violating the assumptions of MSC and loci across our large dataset are practically guaranteed to violate the assumption of a single generating tree (Rannala et al. 2020). Recent work has noted discordance between reconstructed gene trees for individual loci and the tree supported by the same loci when part of a larger supermatrix (Walker et al. 2020).

Overall, it is worth keeping in mind both the issue of model fit in short loci leading to erroneous support, as well as the issues of both model fit and lost signal in concatenated datasets, as it is unclear which approach leads to a more accurate phylogenetic signal inference. We thus conclude that both approaches should be used when interrogating phylogenetic conflicts.

### Heterogeneity of Signal within the Dataset and Impacts of Strong Signal Outliers

Our results suggest that even a large dataset with loci relatively uniformly sampled across gene types can have some heterogeneity of phylogenetic signal, with a group of loci with a strong phylogenetic signal favoring a different topology compared to a group of loci with a weak signal. The selected phylogenetic hypothesis would depend on the proportions of these two groups of loci in a dataset. In this context, we suggest exercising caution with picking loci with the top most strong phylogenetic signal, since the phylogenetic error might be different for such subset and mislead the analysis. For many phylogenomic subsampling methods, loci are selected based on alignments between a handful of reference genomes (Faircloth et al. 2012; Lemmon et al. 2012), and are typically checked to reproduce a well-supported phylogeny as a control measure in the design phase (Faircloth 2017). Our results show that such (non-random) filtering could bias results away from the signal across the genome.

### Signal Metrics Discordance on a Locus-by-locus Basis

Despite the congruence between mean ΔlnL and mean sCF in the entire dataset, there was no strong correlation between the two metrics on a locus-by-locus basis. This is not surprising given that sCF is computed in a parsimony framework and is not impacted by model parameter misspecification. On the other hand, sCF is a quartet-based metric and can be stochastic when the conservative nature of SISRS loci results in a small number of informative quartets (Minh et al. 2020a). Interestingly, there was a relatively strong positive correlation between ΔlnL and sCF for the loci supporting the monophyletic primates. We hypothesize that these loci, even despite their short size, have enough signal to overcome stochasticity in model fit and/or quartet sampling, and thus bring the two metrics to congruence. Consequently, that would mean that in the cases with a strong ΔlnL-sCF signal discordance loci might have elevated levels of noise compared to the phylogenetic signal. We hypothesize that model fit accuracy might explain some of the differences between the two types of ΔlnL-based signal and the sCF-based signal assessments.

### Loci Properties and Signal for Treeshrew Placement

It has been shown that locus type (Literman and Schwartz 2021) and locus properties (Mongiardino Koch 2021) are related to the overall amount of phylogenetic signal. However, we did not observe any large differences in locus properties with respect to a particular conflicting topology (see Figures). While minor but significant differences of some properties could be predictive of the amount of signal and noise in the data, more sensitive methods would be needed to tease this out. It is also possible that the signal for each of the topologies is randomly distributed throughout the genomes independent of the locus properties, and any difference found between groups of loci favoring competing hypotheses would bear no implication across different datasets and phylogenetic problems.

### Taxon Sampling and Signal for Treeshrew Placement

Taxon sampling has been shown previously to affect phylogenetic estimates (Rosenberg and Kumar 2003; Nabhan and Sarkar 2012). In this study, Dermoptera, Lagomorpha, and treeshrews were represented by only a few species each and were constrained to have at least one species present in each locus. Thus the sampling of the diverse Primates and rodents primarily determined the overall variance in taxon sampling, as well as the fractions of Primatomorpha and Glires. While our taxon set was relatively skewed towards a high fraction of Glires (0.55), similar fractions of Glires were included in prior analyses supporting all alternative hypotheses of treeshrew placement. In the datasets recovering the Glires sister, the fraction of Glires ranged between 0.39 and 0.44 (Song et al. 2012; Romiguier et al. 2013; Tarver et al. 2016; Liu et al. 2017), while in the datasets finding the Primatomorpha sister this fraction ranged between 0.31 and 0.5 (Hallström and Janke 2008; McCormack et al. 2012; Kumar et al. 2013; Esselstyn et al. 2017). Thus, although the high Glires sampling might be seen as a concern, our results are consistent with results from lower Glires sampled datasets. However, it is possible that more intricate properties of taxon sampling (such as presence or absence of particular species) are at play here.

### SISRS Data and Conflict Interrogation

Our results also contain several practical outcomes with regard to SISRS-based data and the newly introduced orthologous locus extraction pipeline. The recovered SISRS loci in our case are relatively conserved, which is unsurprising given the methodology of the pipeline. Although SISRS alignments were considerably shorter (∼200bp vs ∼700bp (Esselstyn et al. 2017)]), the proportion of parsimony informative sites in our dataset was in the same range as in some of the mammal UCE studies (McCormack et al. 2012; Esselstyn et al. 2017), amounting on average to about 70 parsimony informative sites per alignment. Small locus size, however, should not impact the results much since we did not use individual loci to produce and analyze gene trees (apart from the BLC filtering). It is conceivable that the model parameter estimation could be compromised when small loci fit to topologies individually. On the other hand, longer loci are likely to violate the single underlying tree assumption. While it is unclear which option balances the phylogenetic signal and noise better, we explored both individual fit and concatenation fit approaches in detail. Additionally, we also used other metrics to estimate phylogenetic signal, including the total likelihood summed across all sites of all loci, thus we argue that short loci produced by SISRS do not pose any significant obstacle to the result interpretation.

Given that SISRS loci contain considerable proportions of both coding and regulatory elements, as well as a large amount of normally not used unannotated (intergenic) loci (Literman and Schwartz 2021), SISRS datasets offer a good diversity of locus types. In contrast, most other methods focus on subsets of particular locus types (i.e., UCE, protein-coding sequences, introns) or small subsets of randomly targeted markers (i.e., RADSeq methods), which have the potential to bias results (Literman and Schwartz 2021).

## Conclusion

Our work highlights the importance of per-locus phylogenetic signal interrogation (Smith et al. 2015; Brown and Thomson 2017; Shen et al. 2017) as a valuable tool to examine support for alternative phylogenetic relationships. While our initial phylogeny estimation supported the treeshrew as the sister to Primatomorpha, this interrogation process revealed a quite pronounced support for the Primatomorpha+Glires sister hypothesis as well. The methodology for conflict interrogation utilized in this study allowed us to ascertain the overall magnitude of the conflict and the amount of signal supporting each conflicting topology in a locus-by-locus framework. The techniques allowed us to disentangle the phylogenetic signal of strong signal outliers from that of the remainder of the dataset and discover that these can contain support for different topologies. A non-random subsetting of strong signal outliers could thus bias results away from the signal across the genome and mislead the phylogenetic inference. This suggests that filtering loci (or selecting particular loci to sequence) should be done with caution due to the potential to estimate a phylogeny that is not concordant with an estimate based on a more complete dataset. Using locus-by-locus interrogation we also found discordance between the model-based and parsimony-based methods, which we in part attribute to the adequacy of model fit to the data.

Overall, with a wealth of phylogenetic information provided by SISRS, we support two alternate phylogenetic placements for treeshrews, even though our initially estimated phylogeny strongly supported a single hypothesis. Two hypotheses, Primatomorpha-sister or Primatomorpha+Glires-sister receive statistical support, depending on the method of phylogenetic signal assessment, as well as whether the signal outlier loci were included or excluded from the analysis. This observation, even with an extremely large DNA sequence dataset, suggests that treeshrews, Glires, and Primatomorpha might have diversified in rapid succession.

## Supporting information

Tab. S1. Sample information

Tab. S2. Loci properties and signal

Tab. S3. Results of the statistical analyses

File. S1. Compressed sequences of SISRS loci

## Acknowledgments

This research was funded by a grant to R. Schwartz from the National Science Foundation (DBI-1942273). We thank members of the Schwartz lab, as well as three reviewers, for valuable comments and suggestions. The High Performance Research Computing facility at URI is acknowledged for providing the computational resources for the analyses in this manuscript.

## Supplementary information

Data available from the Dryad Digital Repository: http://dx.doi.org/10.5061/dryad.[NNNN]

### Supplementary Figures

**Fig. S1.**
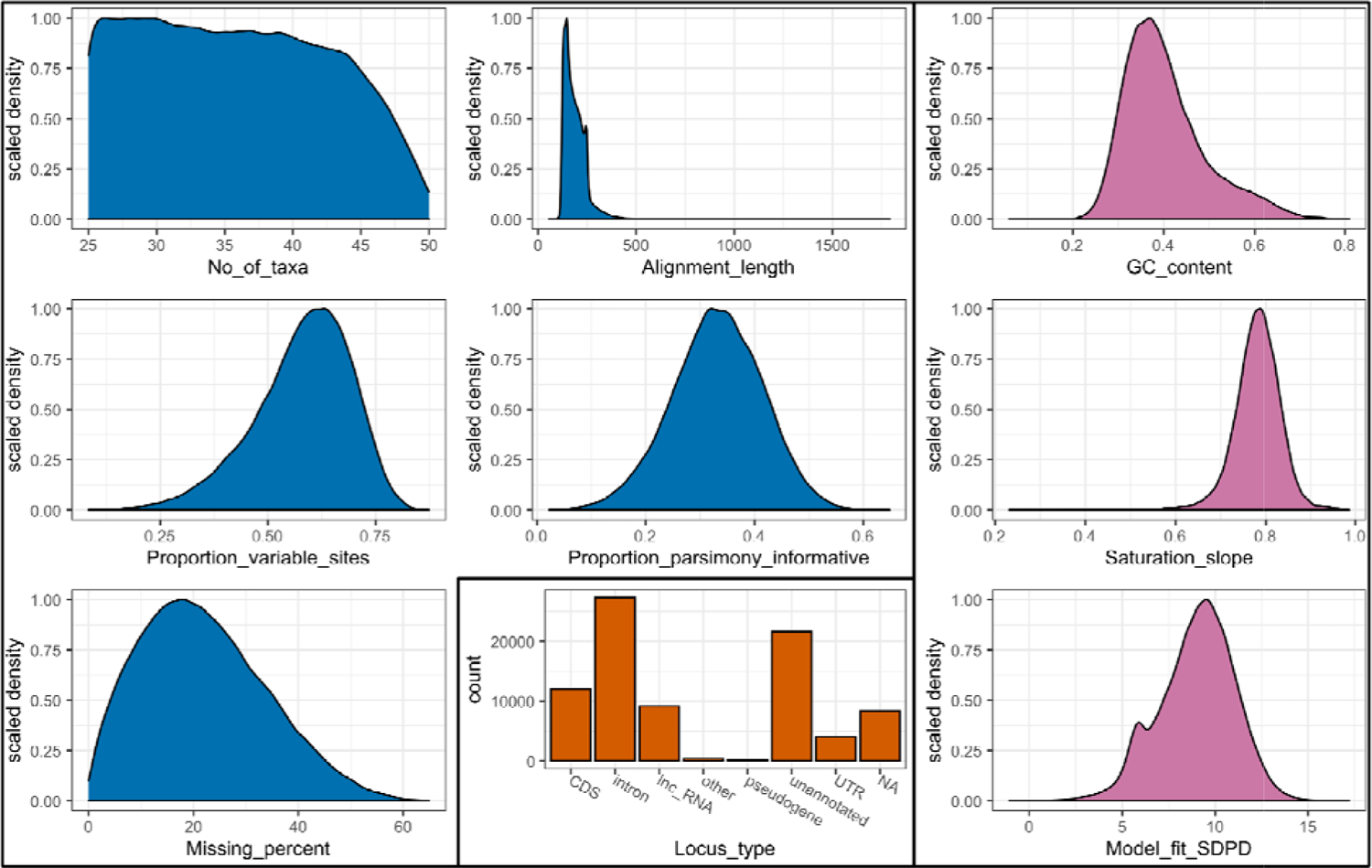
Properties of the evaluated SISRS loci. A set of properties on the left (blue) contains size and information related features, a set of properties on the right (pink) contains compositional features, the bottom center section shows counts of different locus types.

**Fig. S2.**
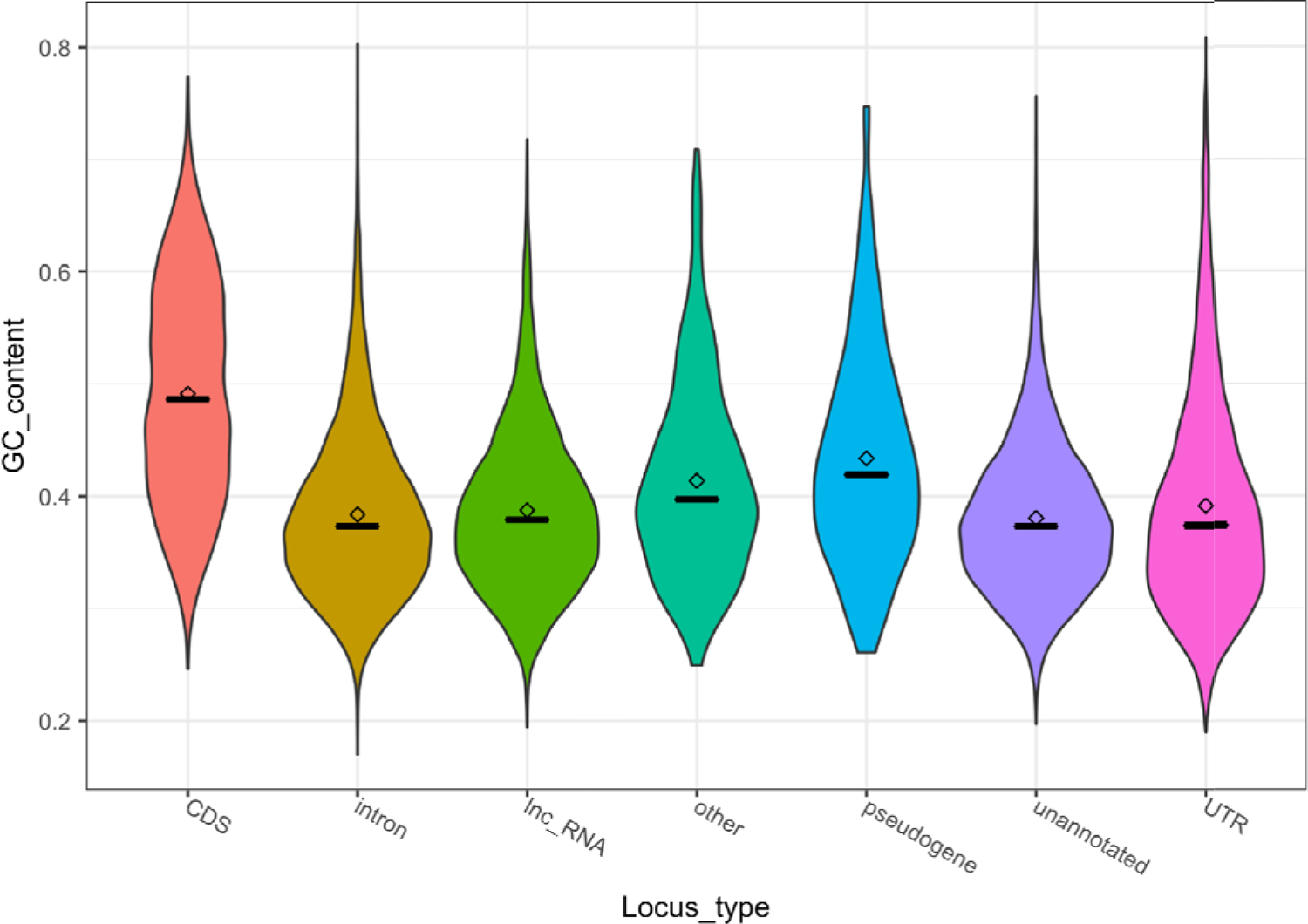
GC content of different locus types in the SISRS dataset.

**Fig. S3.**
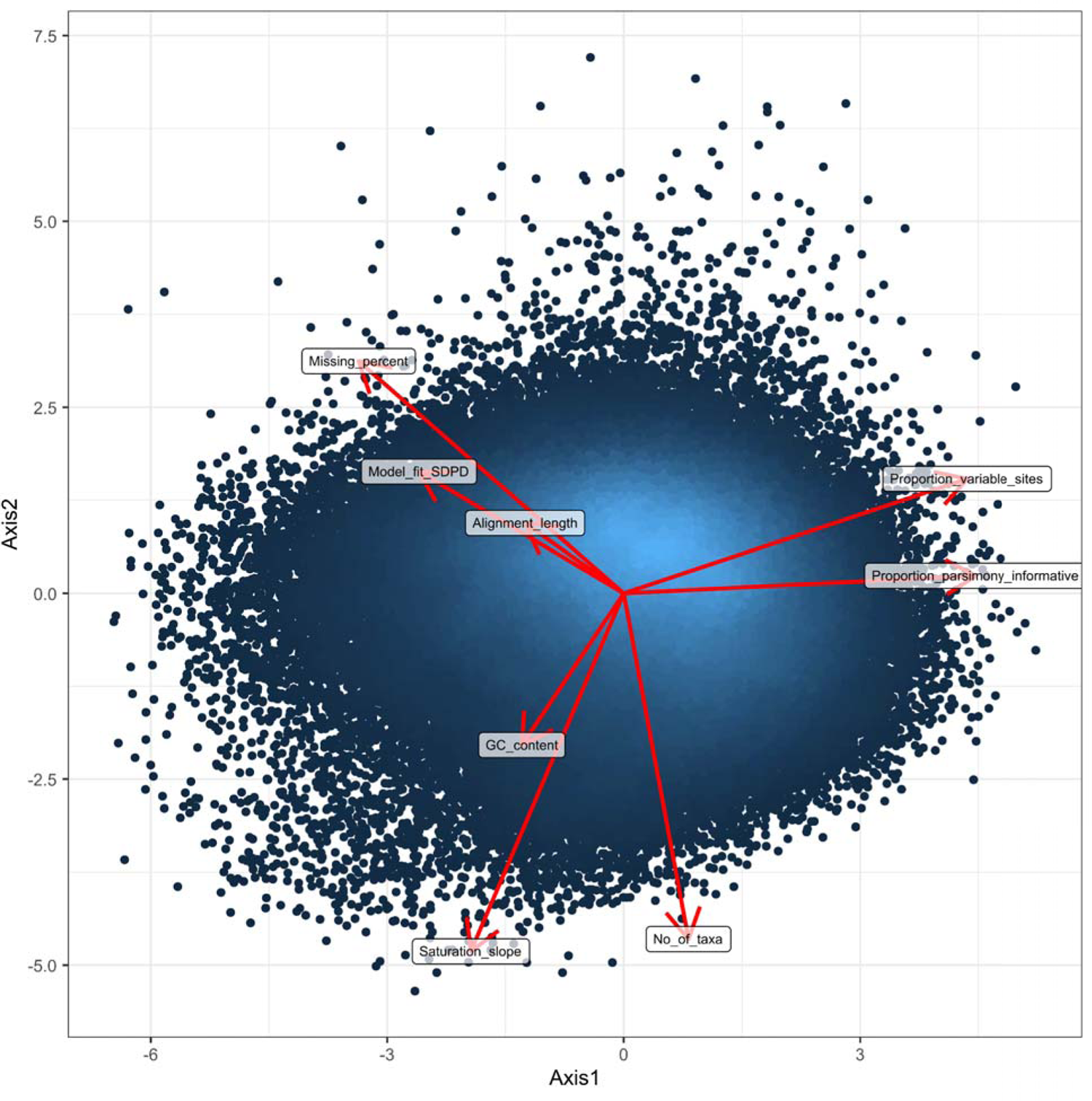
Interrelationship between evaluated loci properties, assessed via a principal component analysis. Only the first two PCA components are shown.

**Fig. S4.**
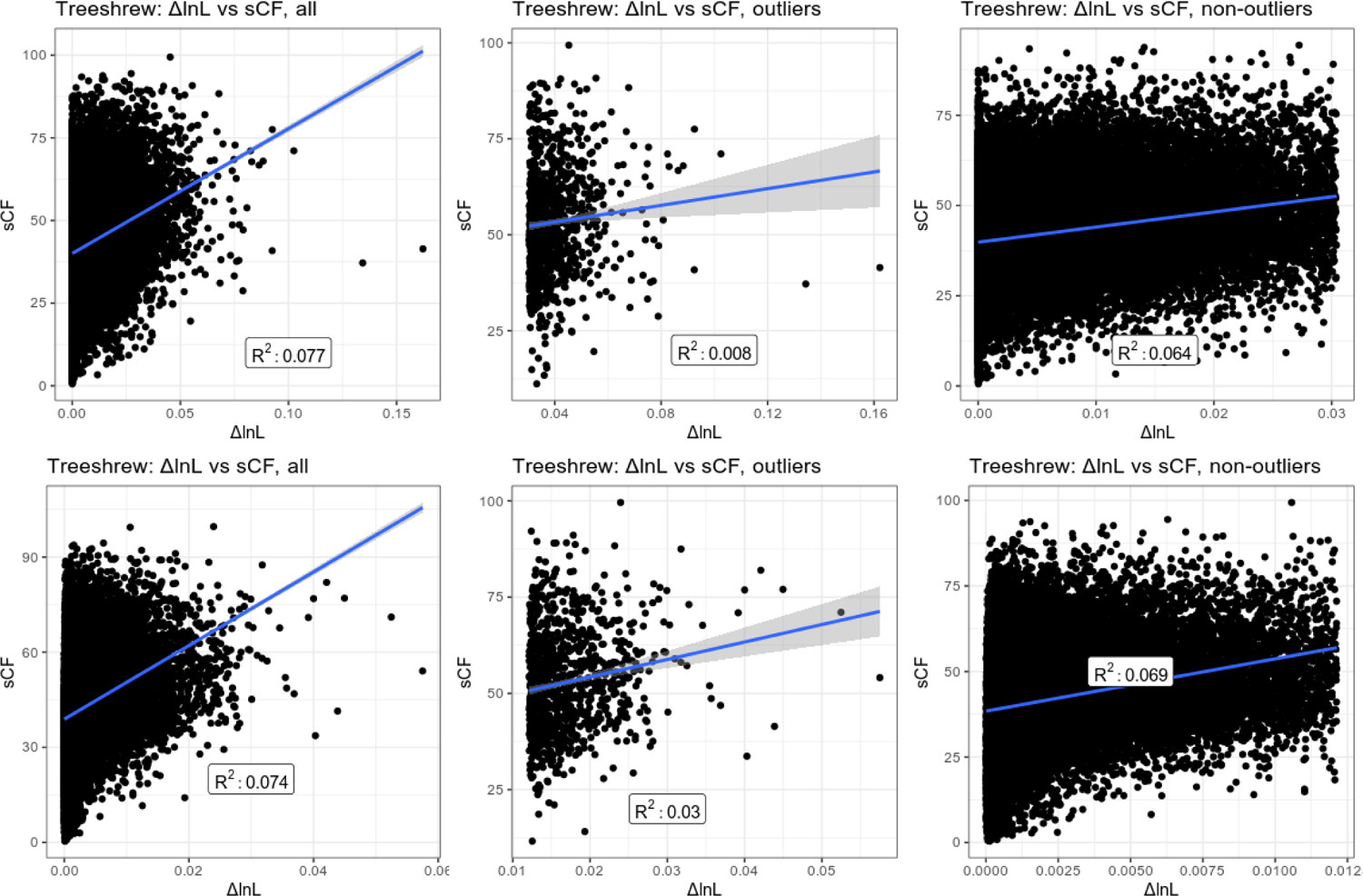
Interrelationship between ΔlnL and sCF with respect to competing treeshrew placements. Top panels correspond to individual locus fit, bottom to fit in concatenation.

**Fig. S5.**
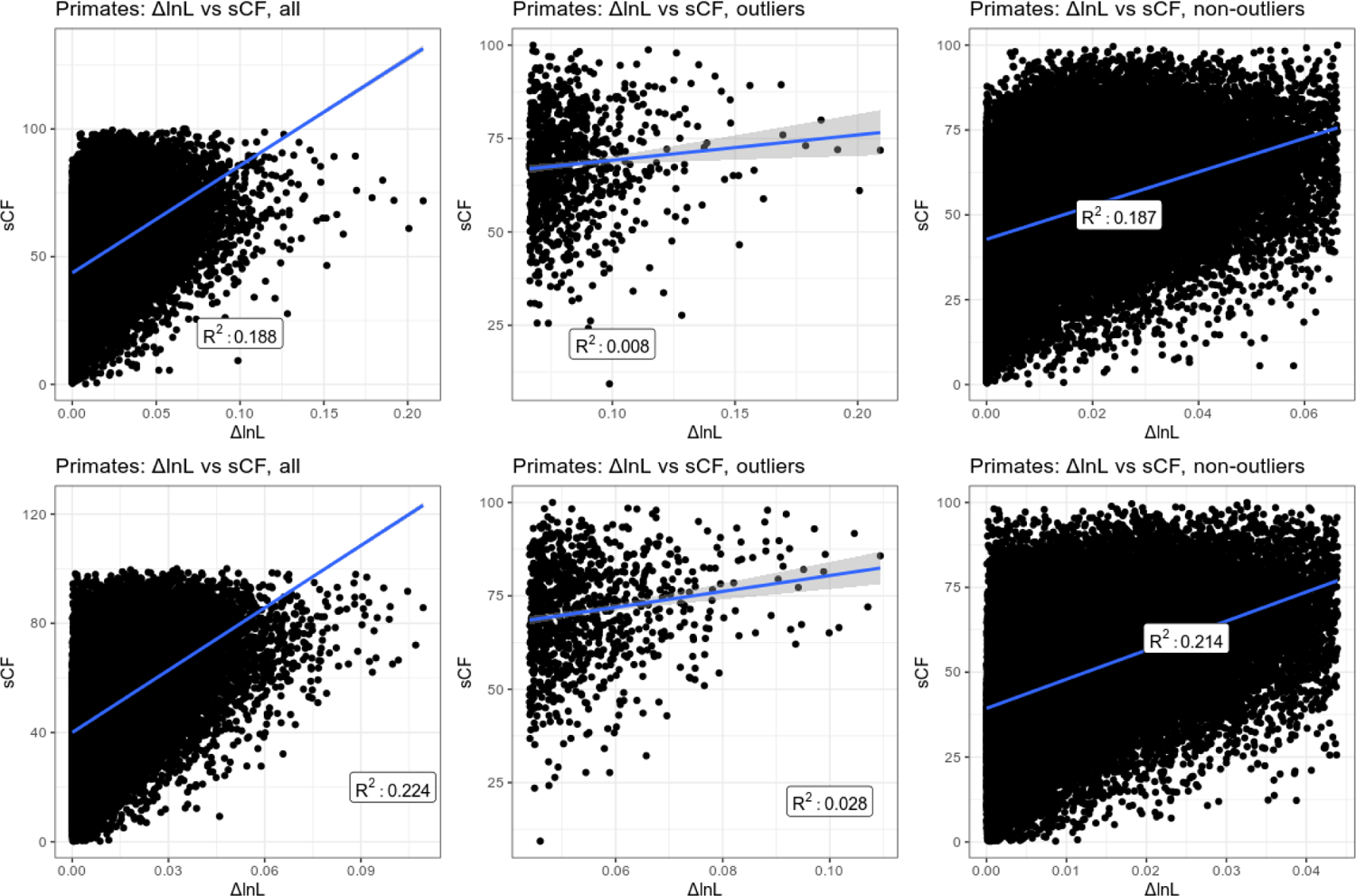
Interrelationship between ΔlnL and sCF with respect to monophyly of primates. Top panels correspond to individual locus fit, bottom to fit in concatenation.

### Supplementary Tables

Tab. S1. Sample information

Tab. S2. Loci properties and signal

Tab. S3. Results of the statistical analyses Miscellaneous

File. S1. Compressed sequences of SISRS loci

